# Decoding the chemical language of *Suillus* fungi: genome mining and untargeted metabolomics uncover terpene chemical diversity

**DOI:** 10.1101/2023.11.20.567897

**Authors:** Sameer Mudbhari, Lotus Lofgren, Manasa R. Appidi, Rytas Vilgalys, Robert L. Hettich, Paul Abraham

**Affiliations:** UT-ORNL Graduate School of Genome Science and Technology, University of Tennessee, Knoxville, TN, USA; Biosciences Division, Oak Ridge National Laboratory, Oak Ridge, TN, USA; Biology Department, Duke University, Durham, NC 27708, USA

**Keywords:** Fungi, ectomycorrhizae, *Suillus*, genome mining, metabolomics, secondary metabolites, terpene

## Abstract

Ectomycorrhizal fungi establish mutually beneficial relationships with trees, trading nutrients for carbon. *Suillus* are ectomycorrhizal fungi that are critical to the health of boreal and temperate forest ecosystems. Comparative genomics has identified a high number of non-ribosomal peptide synthetase and terpene biosynthetic gene clusters (BGC) potentially involved in fungal competition and communication. However, the functionality of these BGCs is not known. This study employed co-culture techniques to activate BGC expression and then used metabolomics to investigate the diversity of metabolic products produced by three *Suillus* species (*S. hirtellus* EM16, *S. decipiens* EM49, and *S. cothurnatus* VC1858), core members of the Pine microbiome. After 28 days of growth on solid media, liquid chromatography–tandem mass spectrometry identified a diverse range of extracellular metabolites (exometabolites) along the interaction zone between *Suillus* co-cultures. Prenol lipids were among the most abundant chemical classes. Out of the 62 unique terpene BGCs predicted by genome mining, 116 putative terpenes were identified across the three *Suillus* species using metabolomics. Notably, some terpenes were significantly more abundant in co-culture conditions. For example, we identified a metabolite matching to isomers isopimaric acid, sandaracopimaric acid, and abietic acid, which can be found in pine resin and play important roles in host defense mechanisms and *Suillus* spore germination. This research highlights the importance of combining genomics and metabolomics to advance our understanding of the chemical diversity underpinning fungal signaling and communication.

**Importance:** Using a combination of genomics and metabolomics, this study’s findings offer new insights into the signaling and communication of *Suillus* fungi, which serve a critical role in forest ecosystems.

## Introduction

Ectomycorrhizal (ECM) fungi are important community members in temperate and boreal forest ecosystems, where they form obligate symbiosis with most woody plant species. Ectomycorrhizal fungi trade fungal-scavenged nutrients such as nitrogen and phosphorus for host-derived photosynthetically fixed carbon and play critical roles in biogeochemical cycling [1]. Fungi in the genus *Suillus* are important ECM mutualists that associate almost exclusively with host trees in the family Pinaceae [2], where they facilitate improved seedling establishment, drought resistance via improved water conductance, and ecological remediation of heavy metal contaminated sites [3-5]. In their role as essential root symbionts, ECM fungi interact with complex consortia of other organisms including plants, bacteria, viruses, invertebrates, and other fungi. Interactions between ECM fungi and these co-occurring community members are mediated by complex chemical signals including small molecules, proteins, and secondary metabolites [6]. Despite the importance of these interactions, our understanding of the chemical diversity facilitating interactions between ECM fungi, their hosts, and the rhizosphere community is limited. Characterizing the identity of the metabolites involved in these interactions, and the conditions under which they are expressed, are the first steps to elucidating the diverse ecological functions of secondary compounds in environmental sensing, competition, and communication.

Unlike primary metabolites, secondary metabolites are not essential to cellular functions but often play critical roles in signaling, communication, and regulating inter- and intra-species interactions between fungi and their surrounding communities [7]. To date, most studies of fungal secondary metabolic diversity have focused on pathogens, saprophytes, and endophytes, particularly those in the phylum Ascomycota [8]. This hinders our ability to appreciate the vast repertoire of secondary metabolites produced by fungi occupying diverse lifestyles. Previous studies of *Suillus* metabolomes have mostly focused on fruit bodies (Mushrooms) [9-11] and were able to identify metabolites such as prenylated phenols and boviquinones [10]. The exometabolomes of *Suillus*, on the other hand, have not been well characterized. Previous work using comparative genomics and genome mining-based predictions of Biosynthetic Gene Clusters (BGCs) in *Suillus* indicated that the genus may have a significantly higher capacity to produce terpenes and non-ribosomal peptides than other ECM species [12]. However, genome-mining-based approaches are unable to characterize most of these compounds past these broad metabolite classes, and the ecological role and conditions necessary for their expression are unknown. Further, genes within BGCs are most often not expressed under standard laboratory conditions [13]. As such, the induction of BGCs requires specific environmental conditions. Techniques such as OSMAC (one strain many compounds) have been successfully used to activate silent BGCs by systematically altering biotic and abiotic environmental variables [14]. Today, it’s widely recognized that one of the most efficient and effective forms of metabolite induction is coculturing fungi with other organisms [15].

Unlike many OSMAC strategies, coculture has the added benefit of being able to address ecologically relevant scenarios, including interactions between naturally co-occurring or co-evolving community members. As a first step towards characterizing the exometabolite chemical diversity in *Suillus*, we chose three genome-sequenced species of *Suillus* known to co-occur and associate with the same species of host tree (*Pinus taeda* – Loblolly Pine). Genome-mining was used to predict and study the similarity between secondary metabolite-producing BGCs across the three species. The three species were grown in monoculture as well as coculture for all pairwise combinations. Untargeted metabolomics was then used to characterize the exometabolites produced at the growth interface between two fungal cultures.

## Materials and Methods

### 2.1. Genome mining

Genomes for the three species of *Suillus* used in this study; *S. decipiens* EM49, *S. cothurnatus* VC1858, and *S. hirtellus* EM16, were first published and characterized in [12] and are publicly available from the JGI MycoCosm database [16]. Biosynthetic Gene Clusters were predicted using antiSMASH v.6.0.1 [17], with the parameters (--taxon fungi --cb-general --cb-subclusters --cb-knownclusters --p fam2go). Orthology and conservation predictions between BGC were carried out via BiG-SCAPE with default parameters [18].

### 2.2. Co-culture and growth assay

The three species of *Suillus* were inoculated onto 100mm petri dishes containing solid high carbon Pachlewski’s media [19]. Each petri dish was inoculated with n=2 four-millimeter plugs placed exactly two centimeters from one another and equidistant to a diameter line intersecting the plate. Treatments included all pairwise combinations of the three *Suillus* species (at n= 5 technical replicates). Single-species controls were inoculated with two plugs as above, and negative controls included uninoculated media from the same batch of plates used for treatment construction. Cultures were grown for 28 days, in the dark, at room temperature. Starting at seven days post inoculation (dpi), colony area was measured twice per week by using background illumination and outlining the colony margins on the bottom side of each petri dish using a fine-tip marker. At the end of the 28-day growth period, we captured images of the bottom of each petri dish using a desktop scanner and calculated colony area at each time point using the program imageJ [20]. After 28 days, 3 agar plugs were collected and placed in 1.5 mL cryotubes using a sterile brass core-borer, taking plugs from along the diameter line of the plate capturing the interaction zone between the two cultures. After collection, samples were immediately frozen in liquid nitrogen, and stored at --80C until sample processing.

### 2.3. Metabolomics sample preparation

The frozen agar plugs containing mycelia were lyophilized using a Labconco Freezone freeze dryer (Labconco Equipment Co., KS, USA) until completely dry. The freeze-dried agar plugs were then processed using a biphasic extraction method by mixing 0.5 mL of cold Liquid chromatography coupled with a mass spectrometer (LC-MS) grade water with 0.5 mL of cold hydrated ethyl acetate, vortexed for 1 min and then kept at 4°C overnight for extraction. The ethyl acetate and water fractions were then separated by aspiration. For the aqueous fraction, samples were filtered using a 10 KDa filter (Sartorius Vivaspin® 2 Centrifugal Concentrator Polyethersulfone) by centrifugation at 4,500 x g to remove remaining agar particulates. After filtering, the aqueous extract was freeze-dried and resuspended in an aqueous solvent (5% acetonitrile, 0.1% formic acid) while the ethyl acetate extract was air dried in a chemical fume hood until dry and then resuspended in an organic solvent (70% acetonitrile, 0.1% formic acid). All samples were stored short-term at 4°C until LC-ESI-MS/MS measurements.

### 2.4. Liquid chromatography-electrospray ionization tandem mass spectrometry (LC–ESI-MS/MS)

All samples were analyzed using ultra-high-pressure-liquid-chromatography coupled with a ThermoFisher Q-Exactive Plus mass spectrometer. For each sample, 10 μL inject was flowed across an in-house made nanospray analytical column (75 μm X 150mm) packed with 1.7 μm C18 Kinetex RP C18 resin (Phenomenex). The mobile phase included Solvent A (95% water, 5% acetonitrile, 0.1% formic acid) and solvent B (70% acetonitrile, 30% water, 0.1% formic acid). The metabolites were separated across a 30-minute linear organic gradient (250 nL/min flow rate) from 5% aqueous solvent (5% ACN, 0.1% FA) to 100% organic solvent (70% ACN, 0.1% FA). All MS data was acquired by Xcalibur software version 4.3. using the top N method where N could be up to 5. Target values for the full scan MS spectra were 3 x 10^6^ charges in the 135 – 2000 m/z range with a maximum injection time of 100 ms. Transient times corresponding to a resolution of 70,000 at m/z 200 were chosen and a 2.0 m/z isolation window and isolation offset of 0.5 m/z were used. Fragmentation of precursor ions was performed by stepped higher-energy C-trap dissociation (HCD) with normalized collision energies of 10, 20, and 40 eVs. MS/MS scans were performed at a resolution of 17,500 at m/z 200 with an ion target value of 1.6 x 10^5^ and a 50 ms maximum injection time. Dynamic exclusion was set to 10 s to avoid oversampling measurements of abundant metabolites. A more detailed listing of the parameters can be found in **Supplemental File**. The analysis of untargeted LC-MS/MS data was performed using Thermo Scientific Compound Discoverer (CD) v3.3.1,, GNPS (Global Natural Products Social Molecular Networking)[21, 22] and Skyline v22.2 [23, 24].

## Results

### Fungal Growth was marginally different between inter- and intraspecies pairings

Colony area increased with incubation time for all the fungal cocultures starting seven days post inoculation to day 28 when samples were collected, indicating that all cultures were actively growing when agar plugs were collected for exometabolomic analysis. Generally, isolates grown in intraspecies pairings reached larger colony areas than isolates grown in interspecies (coculture) pairings, but none of these differences were statistically significant (ANOVA, P> 0.05) (**Supplementary File, Figure S1**).

### Genome mining and analysis of biosynthetic diversity predicts specific types of secondary metabolites

Using antiSMASH v6.0.1, we predicted the total number of BGCs for each of the genomes. *S. cothurnatus* VC1858 had the highest predicted number of putative BGCs (53), whereas *S. hirtellus* EM16 and *S. decipiens* EM49 had a lower number of BGCs (43 and 36 respectively) (**Figure 1A**). In general, the total number of predicted BGCs were related to genome size for all three species. As found previously [12], the number of BGCs with genes for non-ribosomal peptide synthetase-like (NRPS-like) proteins and terpene synthases were much higher in all three genomes compared to BGC containing other classes of backbone enzymes, such as Polyketide synthases (PKSs) or hybrid BGCs with multiple types of backbone enzymes. In total, we found 38 terpene synthase BGCs in *S. cothurnatus* VC1858, 23 in *S. hirtellus* EM16, and 19 in *S. decipiens* EM49. Similarly, we identified 14 NRPS-like genes in *S. hirtellus* EM16, 12 in *S. cothurnatus* VC1858, and 14 in *S. decipiens* **(Figure 1B)**

**Figure 1:**
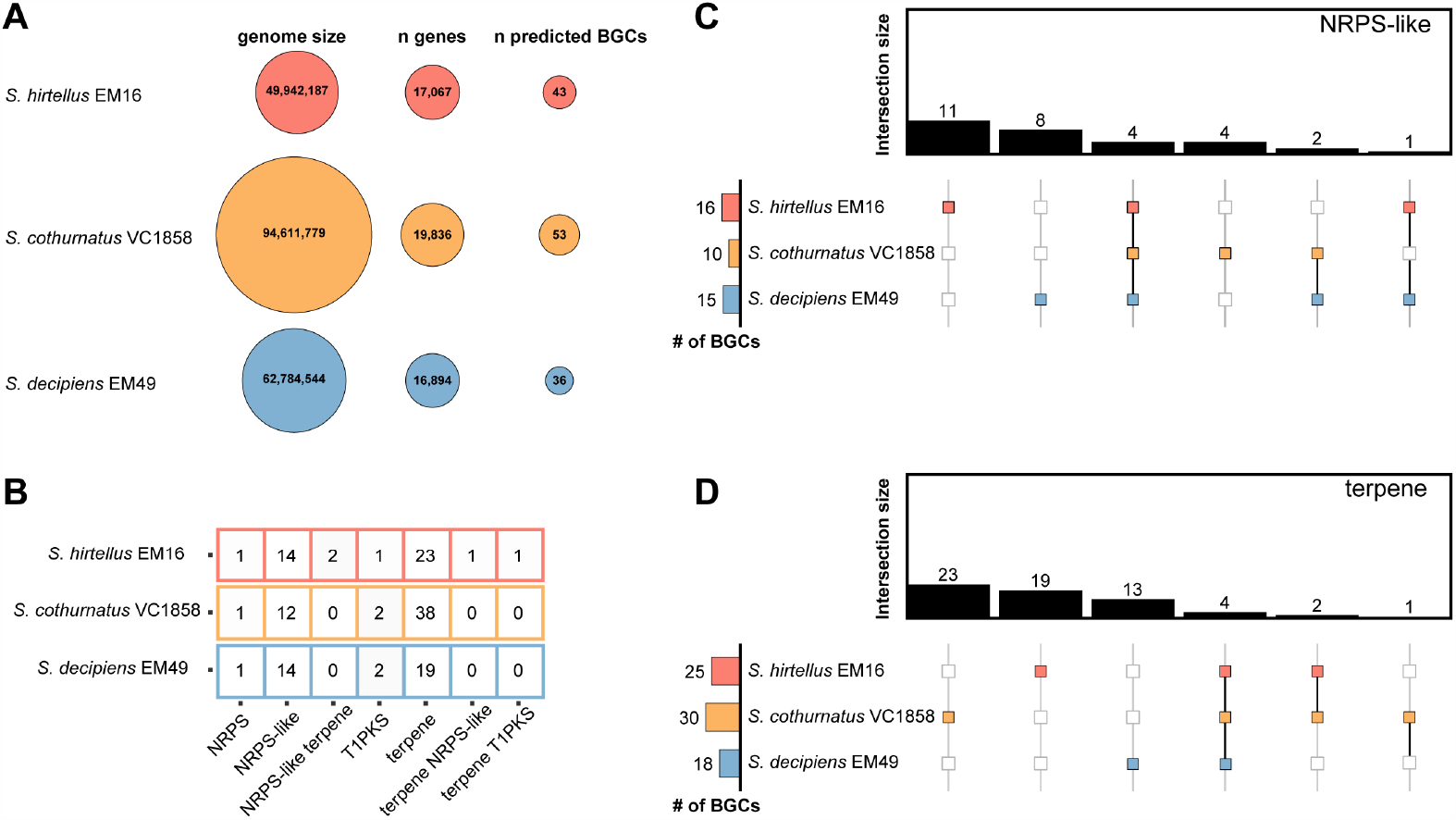
Genome-mining revealed diversity of compounds encoded in three species of *Suillus*. (A) Genome size, number of genes and predicted BGCs varies across *S. cothurnatus* VC 1858, *S. hirtellus* EM16 and *S. decipiens* EM49. (B) The tool antiSMASH v.6.0.1 detected and characterized the number of BGCs in the genome of *S. cothurnatus* VC1858, *S. hirtellus* EM16 and *S. decipiens* EM49. The tool BiG-SCAPE showed low grouping between architecture and sequence similarity for (C) NRPS-like and (D) terpene classified BGCs.

BiG-SCAPE was used to construct sequence similarity networks between BGCs to identify gene cluster families across the three *Suillus* genomes. This analysis revealed that the BGCs containing terpene synthases and NRPS-like genes are remarkably diverse (**Figure 1 C-D**), suggesting there is a vast chemical space to be discovered. Overall, 55 predicted BGCs with terpene synthases did not cluster into gene families based on sequence similarity, 4 were found to be orthologous across the three species of *Suillus*, 2 were orthologous between *S. hirtellus* EM16 and *S. cothurnatus*, and 1 was orthologous between *S. cothurnatus* VC1858 and *S. decipiens* EM49. Regarding BGCs containing NRPS-like genes, 23 did not cluster based on sequence similarity, 4 were orthologous between the three species, 2 were orthologous between *S. cothurnatus* VC1858 and *S. decipiens* EM49, and only 1 orthologous cluster was observed between *S. hirtellus* EM16 and *S. decipiens* EM49.

### Diverse classes of metabolites were identified from the exudates of Suillus during monoculture and coculture conditions

The untargeted metabolomics data processing workflow in Compound Discoverer 3.3 was used for spectral matching. Overall, this workflow predicted a chemical formula for 42,933 compound features that have a measured retention time and a relative abundance based on a calculated peak area (**Supplemental Table 1**). Among these compound features, a metabolite annotation could be assigned to 19,400 features based on accurate precursor mass alone using the ChemSpider database or with further confidence by matching measured fragmentation spectra to spectral libraries in the mzCloud [25] and the high-resolution NIST 2020 [26] database. Next, data was filtered to only consider compound features matching either the mzCloud and/or NIST 2020 databases, and this step resulted in a list of 3,769 putative compound identifications. A principal component analysis was performed separately for the aqueous and organic fraction using MetaboAnalyst version 5.0 [27], and the resulting plots show discrete grouping between biological replicates and separation between coculture groups (**Figure 2 A-B**). Further manual data curation was performed to identify a non-redundant list of compound names that matched with mzCloud and/or NIST 2020 high resolution library and this resulted in a final list of 1,118 putative metabolites observed across this study. ClassyFire version 1.0 [28] was used for chemical taxonomy classification (**Figure 3**), and there were 110 chemical classes observed for 1044 out of 1,118 metabolites (**Supplemental Table 3**). Fatty acyls and carboxylic acids and derivatives were the most abundant chemical classes with 116 metabolites. This was followed by benzene and substituted derivatives with 113 metabolites and prenol lipids with 89 different annotated metabolites. Importantly, prenol lipids encompass 8 different terpenoid subclasses among which terpene lactones, sesquiterpenoids and diterpenoids were the top three most abundant subclasses.

**Figure 2:**
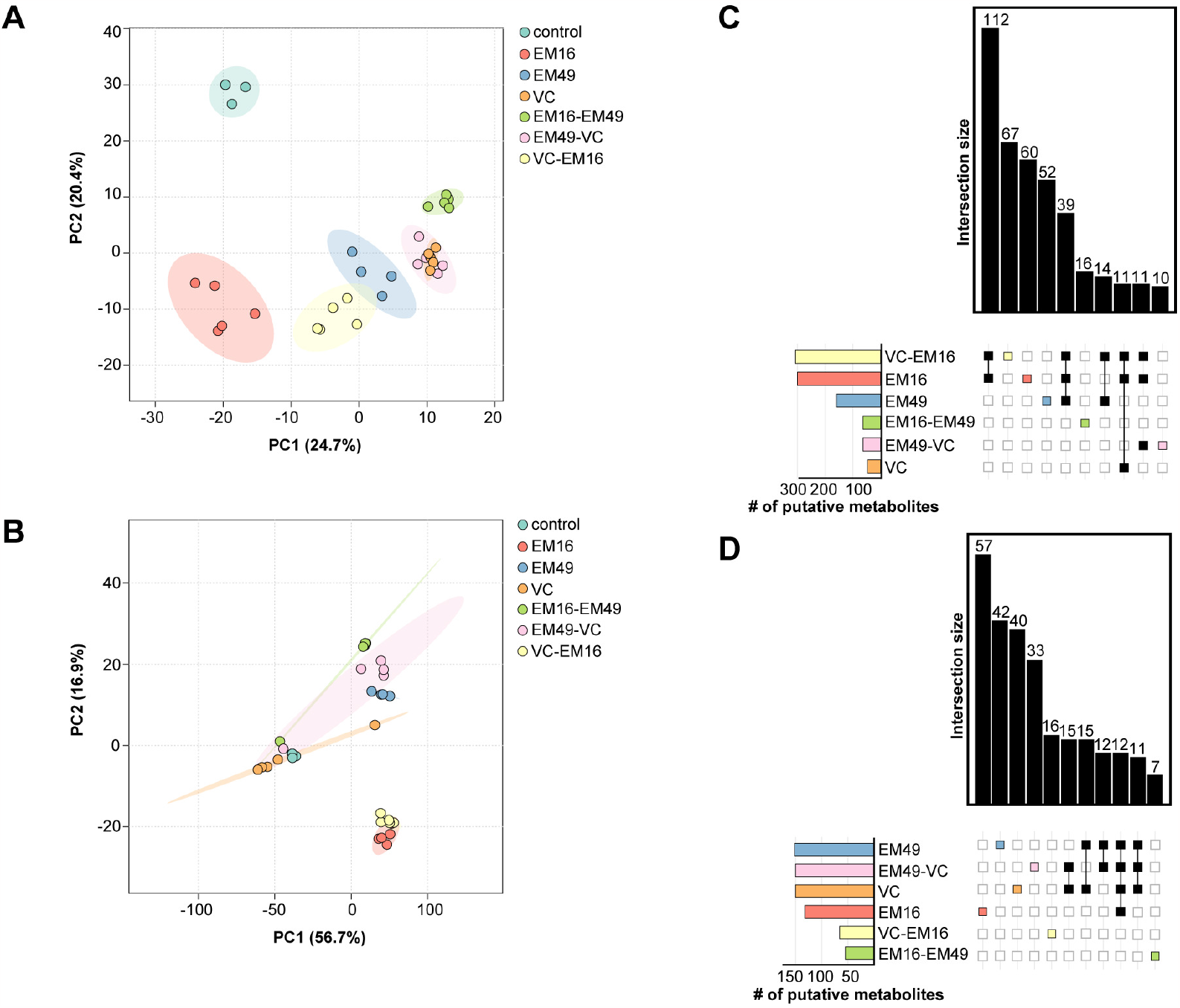
Principal component analysis (PCA) plot and Upset plot showed metabolite variation among different culture conditions. PCA for metabolites observed in (A) ethylacetate fraction and (B) aqueous fraction showed clear separation among culture conditions. The replicates are closer together while culture groups are well separated along the PCA space. Upset plot showing number of putative metabolites that are present in either multiple culture condition or present in only specific culture group for (C) ethylacetate and (D) aqueous fractions.

**Figure 3:**
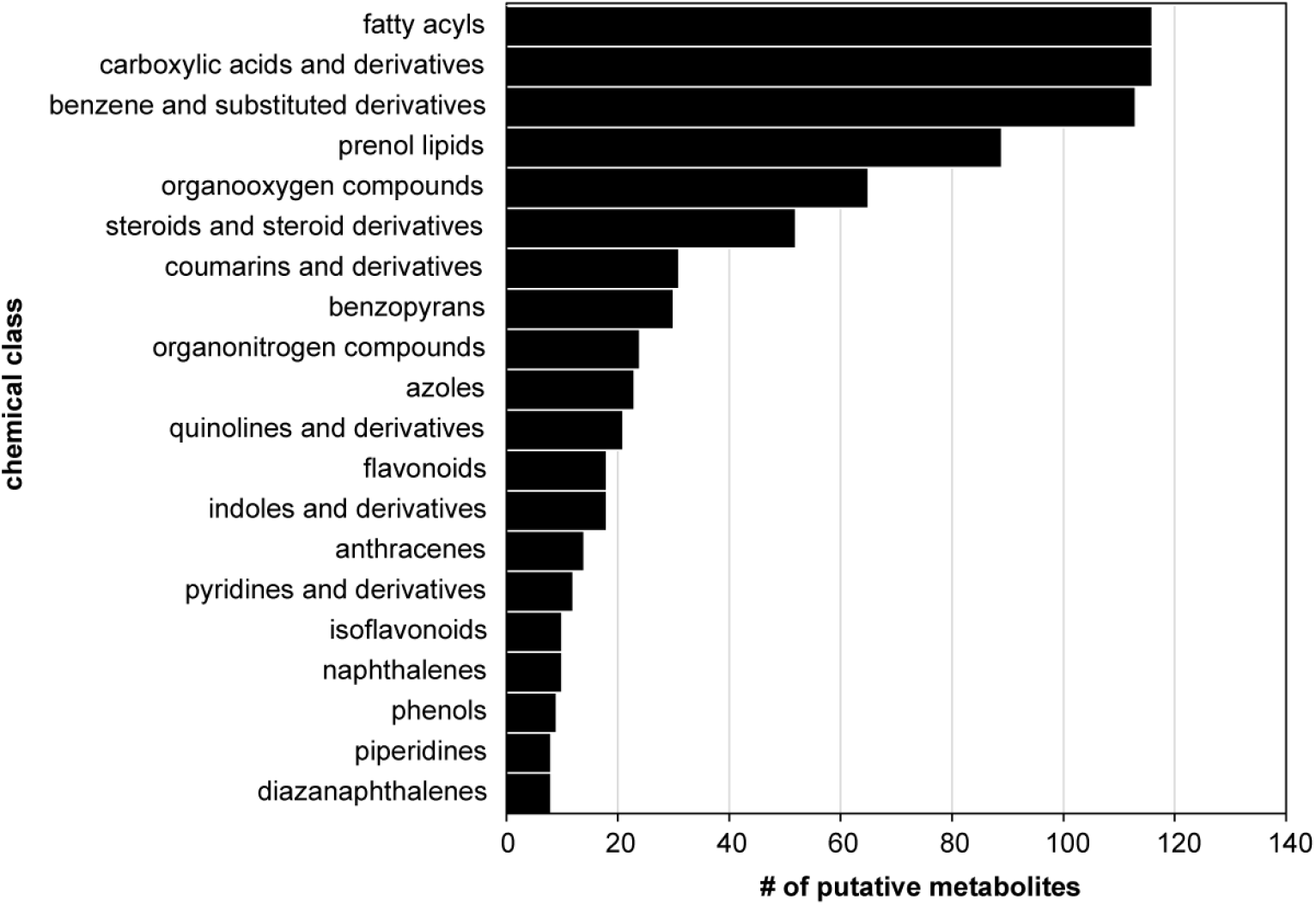
Compound classification highlighted chemical diversity of metabolites. The tool ClassyFire classified the chemical taxonomy of compounds for the compounds identified using Compound discoverer workflow. This analysis showed fatty acids and carboxylic acids and derivatives as the most abundant chemical class observed in LC-MS/MS analysis for the three species of *Suillus* used in this study.

Assessing the degree of overlap in putative identifications observed in the intra- and inter-species cocultures will help define conditions under which a particular metabolite or class of compounds are induced. For this analysis, the data was filtered to remove compound identifications lacking robust quantitative results by only retaining compounds having quality chromatographic peaks detected in at least half the biological replicates of at least one sample group. An UpSet plot was created separately for compound identifications detected in an aqueous or organic fraction sample (**Figure 2 C-D**). In general, both UpSet plots show a relatively low degree of overlap between coculture sample groups. The highest number of unique features were observed among the intraspecies cocultures and S. *hirtellus* EM16 had notably more putative identifications than the other species. Interestingly, analysis of the taxonomical classes shows that the organic fraction of the *S. hirtellus* EM16-*S. decipiens* EM49 pairing had 23 unique metabolites that were absent in monoculture measurements of either species. The most abundant class was the organic organooxygen compounds, prenol lipids and benzopyrans. The organic fraction of the coculture of *S. hirtellus* EM16-*S. cothurnatus* VC1858 had 91 unique metabolites not found in either monoculture, with the most abundant class of identified metabolites also being prenol lipids (10 metabolites). The organic fraction of the coculture of *S. decipiens* EM49-*S. cothurnatus* VC1858 had 36 unique metabolites not found in either monoculture, and the taxonomic classification showed prenol lipids, steroids and steroid derivatives were the most abundant classes of compounds. The organic fraction of the coculture of *S. hirtellus* EM16-*S. cothurnatus* VC1858 resulted in far more unique metabolites compared to all other coculture condition, implicating a responsive interaction between the two fungi that supports expression of many BGCs.

### The exometabolome of Suillus was found to be rich in terpenes which aligns well with genome mining based prediction of terpene coding potential

A relatively large number of putative terpene identifications were observed by untargeted metabolomics based on MS/MS similarity to reference spectra present in the high-resolution NIST 2020 and mzCloud libraries. Overall, 116 different terpenes had a high confidence match to a reference spectra and analysis of taxonomic subclass showed 8 different subclasses of terpenoids out of which terpene lactones, sesquiterpenoids and diterpenoids were the three most abundant ones observed in this dataset (**Supplemental Table 3**).

Overall, most putative terpene identifications were detected regardless of being grown in monoculture or coculture. A subset of terpenes that were relatively abundant in the dataset were further processed by the Skyline software [23, 24] to manually curate metabolite feature extraction for refined quantification. The abundance of these distinct terpenes varied greatly among the culture conditions (**Figure 4**). For example, the metabolite feature matching to the non-volatile diterpenoid sandaracopimaric acid was relatively more abundant in coculture conditions while the sesquiterpenoid, 9,11(13)-Eremophiladien-12-oic acid was relatively more abundant in monoculture of *S. hirtellus* EM16 than when *S. hirtellus* EM16 was cocultured with the other two species. Interestingly, sandaracopimaric acid has been previously quantified in pine resin [29] and seedlings where it was found to play an important ecological role in plant-organism interactions. In an effort to further validate the identification of sandaracopimaric acid in this study, a commercial standard was purchased (A2B Chem, CAS No.: 471-74-9) and measured using the same LC-MS/MS settings. As shown in Figure 4B, there is a high similarity between the tandem mass spectra for the sandaracopimaric acid analytical standard and the *Suillus*-derived metabolite (**Figure 4B**). Previous work studying pine resin and tissue have shown that these chemical fractions also have high concentrations of other non-volatile diterpenoids, including abietic acid, pimaric acid, and isopimaric acid [30-32]. To assess tandem mass spectra fragmentation similarity between the isomers sandaracopimaric and abietic acid, a commercial standard was purchased for abietic acid (Sigma Aldrich, CAS No.: 514-10-3) and measured using the same LC-MS/MS settings. As shown in Figure 4C, the spectral matching between the *Suillus*-derived metabolite and this abietic acid analytical standard is also quite similar. Based on this data alone, it is equally likely that this metabolite is either sandaracopimaric acid or abietic acid.

**Figure 4:**
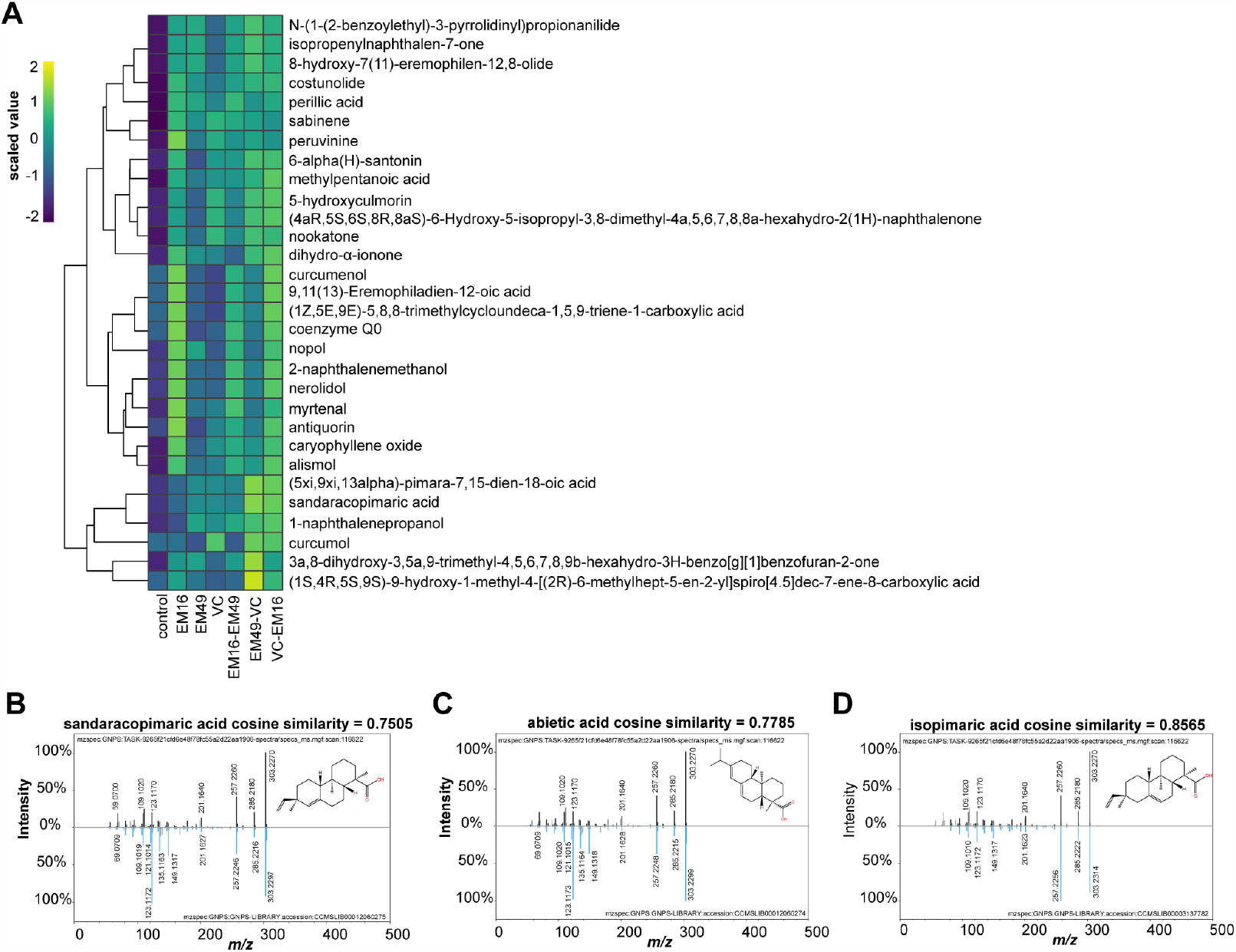
Prenol lipids relative abundance varied among culture conditions. (A) A Heatmap illustrates the relative abundances of selected prenol lipid metabolites that varied in their abundance among culture conditions. These putative identifications were the result of an untargeted metabolomics analysis using Compound discoverer. Mirror match images for (A) sandaracopimaric acid, (B) abietic acid and (C) isopimaric acid. The experimentally observed MS/MS spectrum is shown at the top and a representative MS/MS spectrum from a pure standard is shown in blue color at the bottom.

### Classical molecular networking illustrates Suillus exometabolites diversity and chemical profiles

Molecular networking was used to visualize and explore the entire chemical space beyond what was uncovered by the Compound Discoverer workflow. All experimental data was processed on the GNPS analysis platform (https://gnps.ucsd.edu/; accessed June 28^th^, 2023) in a single workflow using default settings to create a classical molecular network that connects experimental MS/MS spectra (nodes) using a cosine scoring scheme (edges) ranging from 0 (totally dissimilar) to 1 (completely identical). The raw .mzML files from the study were separated into 5 different groups when performing classical molecular networking. All the monoculture files were grouped as one, all three coculture combinations were separately grouped, all media-only controls were grouped together thus resulting in 5 different groups. Experimental MS/MS spectra were matched against the GNPS-community spectral library—a relatively large collection of publicly accessible natural product and metabolomics MS/MS data—to assign putative annotations and identify molecular families, which are related MS/MS spectra differing by simple structural or chemical transformations. The MolNetEnhancer workflow [33] in GNPS was used to combine the outputs from molecular networking and the automated chemical classification through ClassyFire to provide a more comprehensive chemical overview (**Supplementary Table 4)**. With the applied settings, the molecular network contained 16,904 nodes with ∽14% associated with a chemical classification (**Figure 5A**). Among all the nodes, ∽68% of nodes belonged to subnetworks with at least two MS/MS spectra. Analysis of all 16,904 nodes showed that a total of 2,925 nodes were unique to monoculture conditions, 2,482 nodes were unique to coculture conditions, and 1,780 nodes were observed in all the culture conditions yet absent in media blanks. Across the entire study, GNPS provided a putative annotation for 56 total metabolites which were representative of 9 chemical SuperClasses. In general, the top 20 chemical classes and their representation observed by GNPS were similar to what was found using the Compound Discoverer workflow (**Figure 5B**). Further inspection of the resulting prenol lipid annotations revealed that the sandaracopimaric acid-related MS/MS spectra (**Figure 4B**) from the Compound Discoverer workflow matched with high confidence to the stereoisomer isopamaric acid (**Figure 4C**) in the GNPS public library.

**Figure 5:**
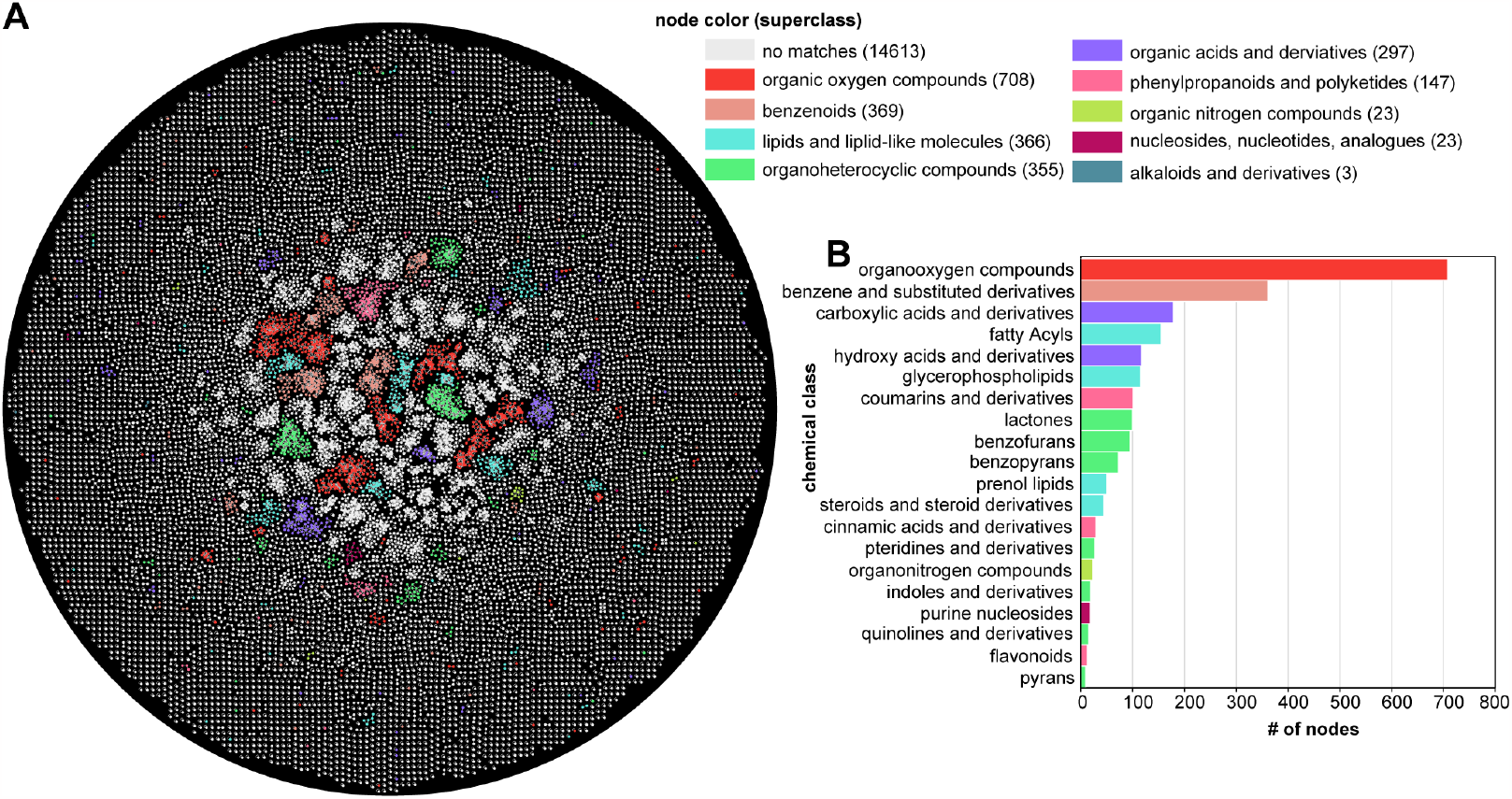
Classical molecular network uncovered predominant annotated and unannotated MS/MS spectra. (A) The GNPS-derived molecular network was visualized by the Cytoscape software. Each node represents a MS/MS spectrum from this study. Nodes colored white represent unannotated MS/MS spectrum and colored nodes represent a MS/MS spectrum associated with a putative metabolite annotation with a relatively high spectral similarity score (correlation >0.7). Nodes with putative annotations were annotated by the MolNetEnhancer workflow in GNPS to illustrate the different superclass annotations. (B) A bar plot shows the number of metabolites belonging to a chemical class reported by the MolNetEnhancer workflow in GNPS.

## Discussion

Pine trees are native to the northern hemisphere, but have been introduced globally for shade, shelter, and wood products, making them some of the most ecologically and economically significant tree groups. In boreal forests, which are typically nutrient poor, Pine nutrition is largely supported by the association with ECM fungi such as *Suillus* species. The establishment, maintenance, and outcome of these ecological interactions depend on a combination of factors, including effective communication between the host and fungal partner. Secondary metabolites are well-known for their role in communication between and within species. When secondary metabolites are released into the extracellular environment (exometabolites), they can convey information about an organism’s presence, function, or metabolic status. These signals help organisms recognize resources and threats. Often, fungal secondary metabolites are not constitutively expressed, and require specific metabolic or environmental cues for induction-complicating the characterization of these important chemical signals.

Previous whole-genome analysis suggested that *Suillus* species have a relatively large number of terpene and NRPS-like BGCs when compared to other ECM fungi [12]. To further assess these highly specialized genomes, we used genome mining and orthology analysis to predict and compare BGCs in three *Suillus* species (*S. hirtellus* EM16, *S. decipiens* EM49, and *S. cothurnatus* VC1858). We found that these three species encoded an abundance of BGCs, which were dominated by species-specific clusters that displayed little conservation between strains. In agreement with previous predictions made across the genus [12], the majority of these clusters were composed of BGCs containing terpene and NRPS-like backbones.

Next, we employed metabolomics to characterize the extracellular secondary metabolites produced by *Suillus* under coculture and monoculture conditions. Conducting a global assessment of metabolites is a significant challenge because variation in sample preparation and metabolite detection introduce biases into the types of chemicals being measured. This motivated us to perform a biphasic extraction to better sample and measure polar and nonpolar fractions of secondary metabolites. It is important to note that no single analytical platform, like LC-MS, is suitable for all metabolomic studies and the selection of which platform to use should be guided by the research question and the nature of the metabolites of interest [34, 35]. For this study, LC-MS was selected for untargeted metabolomic assessment because this method offers high sensitivity, selectivity, and comprehensive coverage for a diverse set of metabolites with different chemical properties [36, 37]. Empowered by the growing availability of LC-MS public data resources, we sought to leverage several available spectral libraries to address another challenge associated with untargeted metabolomics— metabolite identification. In this study, secondary metabolites produced under coculture conditions were mostly fatty acyls, carboxylic acids and derivatives, benzene and substituted derivatives, prenol lipids, and organic oxygen compounds. As expected, only a small amount of the total data (<20%) could be assigned a putative annotation based on spectral matching against a public reference library. While these annotations are useful to describe the physiochemistry of the secondary metabolites observed, it is important to further note that these are putative identifications that require additional verification. The use of commercial standards is recommended to achieve a higher level of confidence. However, as shown for tandem mass spectra matching to several isomers (sandaracopimaric acid, isopamaric acid, and abietic acid), other orthogonal techniques (e.g., nuclear magnetic resonance spectroscopy) must be used to complete the identification process. Nevertheless, the high mass accuracy LC-MS putative identifications presented in this study revealed a vast yet to be characterized landscape of unknown compounds, while providing new insights into the chemical ecology of *Suillus* fungi.

The exometabolome of *Suillus* was found to be rich in terpenes and this observation aligned well with the derived genome mining predictions. Given the ecological importance of terpenes for pine growth and defense, we manually curated quantitative abundances for a subset of putative terpene identifications. Among these, we further interrogated tandem mass spectra belonging to the non-volatile diterpenoid sandaracopimaric acid through an assessment against several available analytical standards. Because the spectral matching between stereoisomer and isomer candidates cannot be differentiated, additional experiments are needed to complete the process of identification. Nevertheless, the prospect of *Suillus* fungi producing either of these compounds is intriguing considering that both are typically produced by conifers such as Pines (John et al., 1993).

While the ecological role of terpene production in ECM fungi is largely unknown, recent metatranscriptomic analysis of pine roots inoculated with different *Suillus* species revealed that terpene synthase genes were differentially expressed during incompatible-host parings (Liao et al., 2016), suggesting that they may play a role in recognition and stress response. The identification of diterpene acids in our *Suillus* cocultures raises several new questions about the ecological roles and origins of terpenes in ECM fungi. Diterpene acids are known to have diverse functions in fungal community interactions; abietic acid is a well-known elicitor of spore germination in *Suillus* (Fries, 1988; Fries, 1990), and has been shown to be antifungal against conifer pathogens such as *Heterobasidion* and *Ophiostoma* [38, 39]. While diterpene acid production is not unheard of in fungi [40], the production of these compounds is typically associated with plants – particularly conifers, raising questions about the origin of diterpene acid BCGs in *Suillus*. Identifying the exact BGCs responsible for encoding these secondary metabolites is nontrivial, but further work to link these products to genes and determine the origin of diterpene acid production in *Suillus* (including the potential for horizonal gene transfer from the host) is warranted.

In conclusion, genome mining coupled with co-culture and untargeted metabolomics revealed a diverse set of secondary metabolites likely to be important for ECM community interactions. In agreement with previous studies, we identified a large number of terpene BGCs using genome mining, encompassing 62 unique clusters. However, this number is relatively small compared to the metabolomic diversity actually produced by this group, which included 116 unique terpenes. As a single BGCs can produce multiple products [41], this incongruence is somewhat expected. However, the staggering difference between the number of BGCs predicted and the number of products identified implies that other factors may be limiting BGC prediction capacity in *Suillus*. Prediction algorithms are typically trained on characterized input data, and because characterized BGCs are underrepresented and understudied in basidiomycetes (particularly in ECM basidiomycetes), prediction algorithms may be more likely to miss BGCs that differ significantly from the training set in structure or sequence similarity. Importantly, while our coculture BGC induction method proved highly effective in producing a variety of terpenes, these within-genus pairings represent only a single type of environmental trigger, and the true terpene diversity of *Suillus* is likely even greater than what we have reported here. Taken together, our LC-MS/MS based untargeted metabolomics analysis of the *Suillus* exometabolome revealed diverse terpenes exuded from species in monoculture and coculture specific conditions highlighting the potential role these chemicals could play in inter- and intraspecific community interfacing.

## Data availability

The data from LC-ESI-MS/MS used in this study are publicly available online at MassIVE (https://massive.ucsd.edu/) with the assigned accession MSV000092795. Similarly, GNPS based classical molecular network result file can be accessed at https://gnps.ucsd.edu/ProteoSAFe/status.jsp?task=fe2b002ce423463da78e55d8adde536f#.

## Acknowledgements

This research is supported the Plant–Microbe Interfaces (https://pmiweb.ornl.gov/) Scientific Focus Areas at ORNL and by a subaward DE-SC0020403 to RV through the U.S. Department of Energy’s Office of Biological and Environmental Research. LL is supported by funding from the National Institutes of Health grant no. T32-AI052080 via the Tri-I MMPTP Fellowship. SM is supported by funding from the Secure Ecosystem Engineering and Design (https://seed-sfa.ornl.gov/) Scientific Focus Area funded by the Genomic Science Program of the U.S. Department of Energy, Office of Science, Office of Biological and Environmental Research (BER) as part of the Secure Biosystems Design Science Focus Area (SFA)

## Figure legends

**Supplementary Figure S1:** Growth differences observed for Suillus species grown in intra- and inter-species pairings. (A, B, C) showing increase in colony area (mm^2^) from Day 0 until Day 28 when agar plugs were taken for exometabolomic analysis for *S. hirtellus* EM16, *S. cothurnatus* VC1858 and *S. decipiens* EM49 respectively when they are grown in either monoculture or in coculture with other species. As can be seen in the figure there is only a marginal growth difference when grown either as a inter species pair or in an intra species monoculture.

